# An Innovative antibody-based Plug-and-Play strategy for SARS-CoV-2

**DOI:** 10.1101/2021.03.11.434589

**Authors:** Gururaj Shivange, Debananda Gogoi, Jogender Tushir-Singh

## Abstract

The novel and highly pathogenic coronavirus (SARS-CoV-2) remains a public health threat worldwide. SARS-CoV-2 enters human host lung cells via its spike protein binding to angiotensin-converting enzyme 2 (ACE2) in a process critical dependent on host protease-mediated fusion event. Thus, effective targeted therapies blocking the first step of viral fusion and cellular entry remains a critical unmet medical need to overcome disease pathology. Here we engineered and describe an antibody-based novel and targeted plug-and-play strategy, which directly competes with the proteolytic activation function of SAR-CoV-2 spike protein. The described strategy involves the engineering of furin substrate residues in IgG1 Fc-extended flexible region of spike targeting antibody. Our results with spike receptor-binding domain (RBD) targeting CR3022 antibody support blockade of the viral function using proof of concept ACE2 overexpressing cells. Our study reveals analytical, safe, and selective mechanistic insights for SARS-CoV-2 therapeutic design and is broadly applicable to the future coronaviridae family members (including mutant variants) exploiting the host protease system for cellular entry.

---

Despite remarkable speed of vaccine approvals, the detailed mechanism of SAR-CoV-2 infectivity and pathology remains unclear(1). SAR-CoV-2 entry into target lung cells is dependent spike interactions with ACE2(2). SAR-CoV-2 spike harbors an arginine-rich multibasic site between attachment (S1) and fusion (S2) domains (Fig. 1A, B), whose cleavage by host cellular furin protease is key for its efficient cellular entry, and cytopathic effects(3). However, host furin protease targeting inhibitors are not specific to primary lung cells infected with SAR-CoV-2(4). Thus, if tested clinically, due to nonspecific tissues distribution, these protease targeting drugs are highly likely to interfere with the normal cellular processes in humans. To counter nonspecificity against SARS-CoV-2 and potential next generation of deadly coronaviridae family exploiting host protease system for cellular entry, here we describe a simple, compelling and targeted approach with the potential therapeutic application.

**Figure 1.**
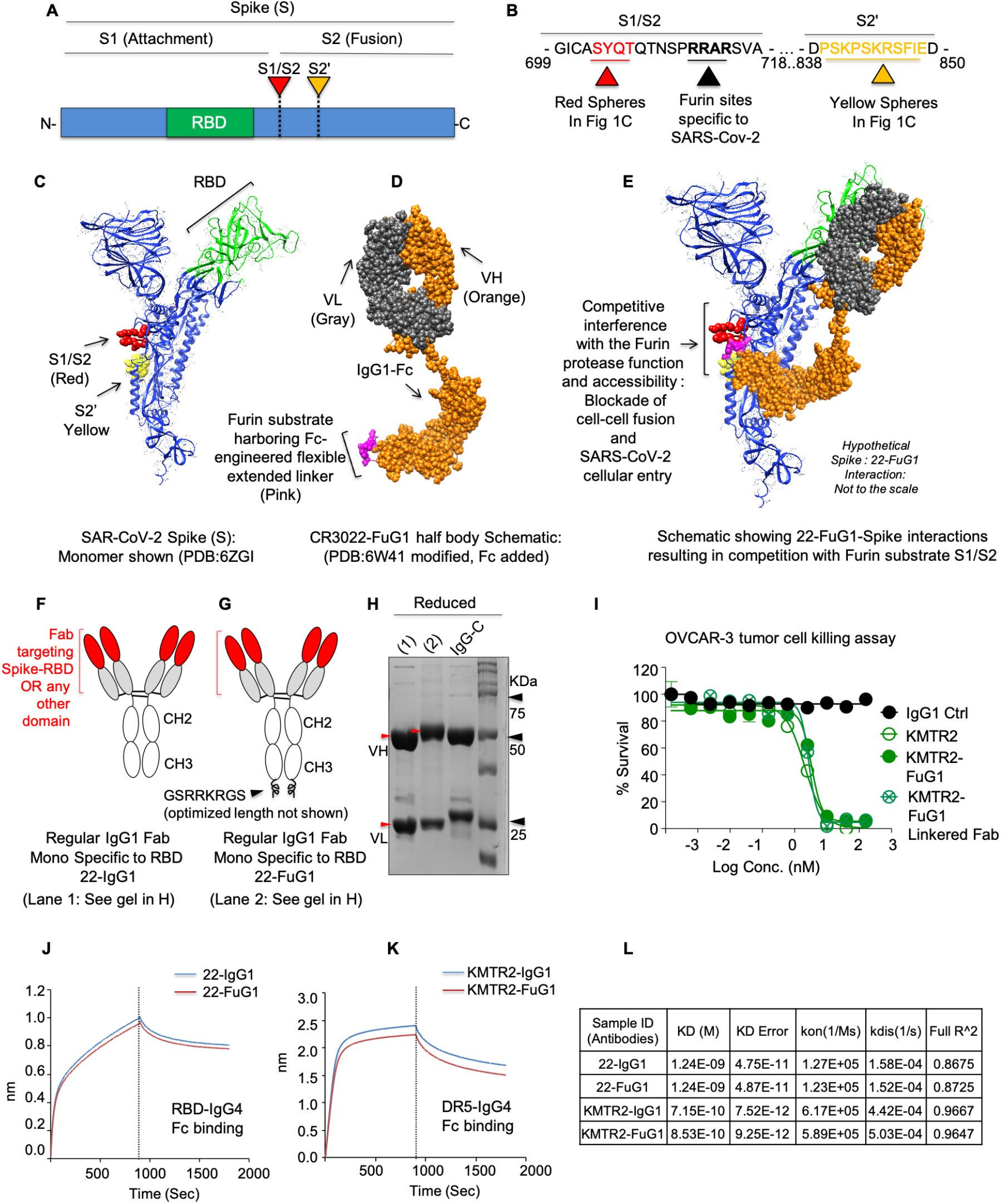
Engineering of FuG1 strategy. (A-B) Schematic and sequence of full spike (S) showing RBD, S1/S2 and S2’ sites. (C) Ribbon structure of SARS-CoV-2 spike monomer (PDB: 6ZGI). Red spheres: S1/S2 residues near furin cleavage site. Yellow spheres: S2’ residues. RBD domain is in green. (D) 22-IgG1 (PDB: 6W41) half-body schematic inserted with Fc extendable linkers harboring furin cleavage residues (Pink). (E) Schematic overlay of 22-FuG1 with SARS-CoV-2 spike monomer (PDB: 6ZGI) depicting competitive interference with furin function. (F-G) Genetic construction schematic of 22-IgG1, 22-FuG1. Black arrow: Fc extended furin linker. (H) A reducing SDS-Gel of 22-IgG1 (1) and 22-FuG1 (2). Red arrows depict ~5 KDa extra size of 22-FuG1 VH. (I) Cell viability assay of KMTR2 and KMTR2-FuG1. (J-L) The binding kinetics of immobilized biotinylated recombinant RBD and DR5-IgG4-Fc against indicated antibodies were measured using BLI.

## Results

We hypothesize that a strategy capable of co-targeting SARS-CoV-2 and furin near the viral infection site is key for board therapeutic specificity. Thus, we engineered an antibody-based strategy that would selectively compete with cellular furin protease function at the site of viral entry. This will inhibit cell fusion ready S2 fragment generation(2, 5) to block cellular infection (Fig. 1C-E). We made use of SARS-CoV-2 RBD-targeting CR3022 antibody (22-IgG1) and reengineered it to harbor optimal furin substrate sites in the Fc-extended linker region (Fig. 1D). We name this approach FuG1, Furin-IgG1 (Fig. 1D-E). FuG1 can be engineered as monospecific (Fig 1F-H) and a linkered bispecific [as described BaCa-2 here(6)] to enhance avidity optimized specificity by co-targeting human lung cell antigens along with spike. The key elements of FuG1 include Fc-extended flexible linkers harboring furin targetable sites (Fig. 1D,G). In reducing-gel conditions, FuG1 antibody’s heavy chain (VH) has ~5 KDa higher molecular weight while the light chain (VL) remains of the same size (Fig. 1H, red arrows). Addition of flexible Fc-extended furin linkers did not affect the function and binding of various FuG1 antibodies (Fig. 1I-L).

When tested, death-receptor-5 (DR5) targeting KMTR2-IgG1 and KMTR2-FuG1 pull-down DR5, however only KMTR2-FuG1 pull-down furin (Fig. 2A-C). Similar results were seen with the 22-FuG1 antibody against spike transfected cells(7) (Fig. 2C). A nonspecific G4S linker without furin targetable residues (22-FuG1-Mut) did not pulldown furin. As 293-ACE2 cells express folate receptor (FOLR1) levels similar to human cancer cells, we also engineered anti-FOLR1, farletuzumab into FuG1. After 2 hours of 293-ACE2 cells transfection with spike, farletuzumab-FuG1 and 22-FuG1 were added. Both FuG1 antibodies were equally effective in inhibiting syncytia, while 22-IgG1 needed significantly higher concentration (Fig. 2D). As FOLR1 is ubiquitous on surface, farletuzumab-FuG1 works by surface occupancy of FOLR1^+^ cells to saturate furin, while 22-FuG1 specifically targets furin after delayed spike expression on cells. In support, delayed treatment after syncytia formation, farletuzumab-FuG1 was significantly ineffective compared to 22-FuG1 (Fig. 2E-G). As expected, FuG1 inhibited spike conversion into S2 in a dose dependent manner. GAPDH revival in lysates indicate reversal of cytopathic cellular state (Fig 2H, I). When tested using spike pseudovirus, 22-FuG1 also inhibited proteolytic activation of spike to generate S2 upon addition on cells (Fig 2J, K). Similar results of syncytia blockade and reduced surface S2 generation were seen in VERO-E6 cells with FuG1 strategy (Fig. 2L-N).

**Figure 2.**
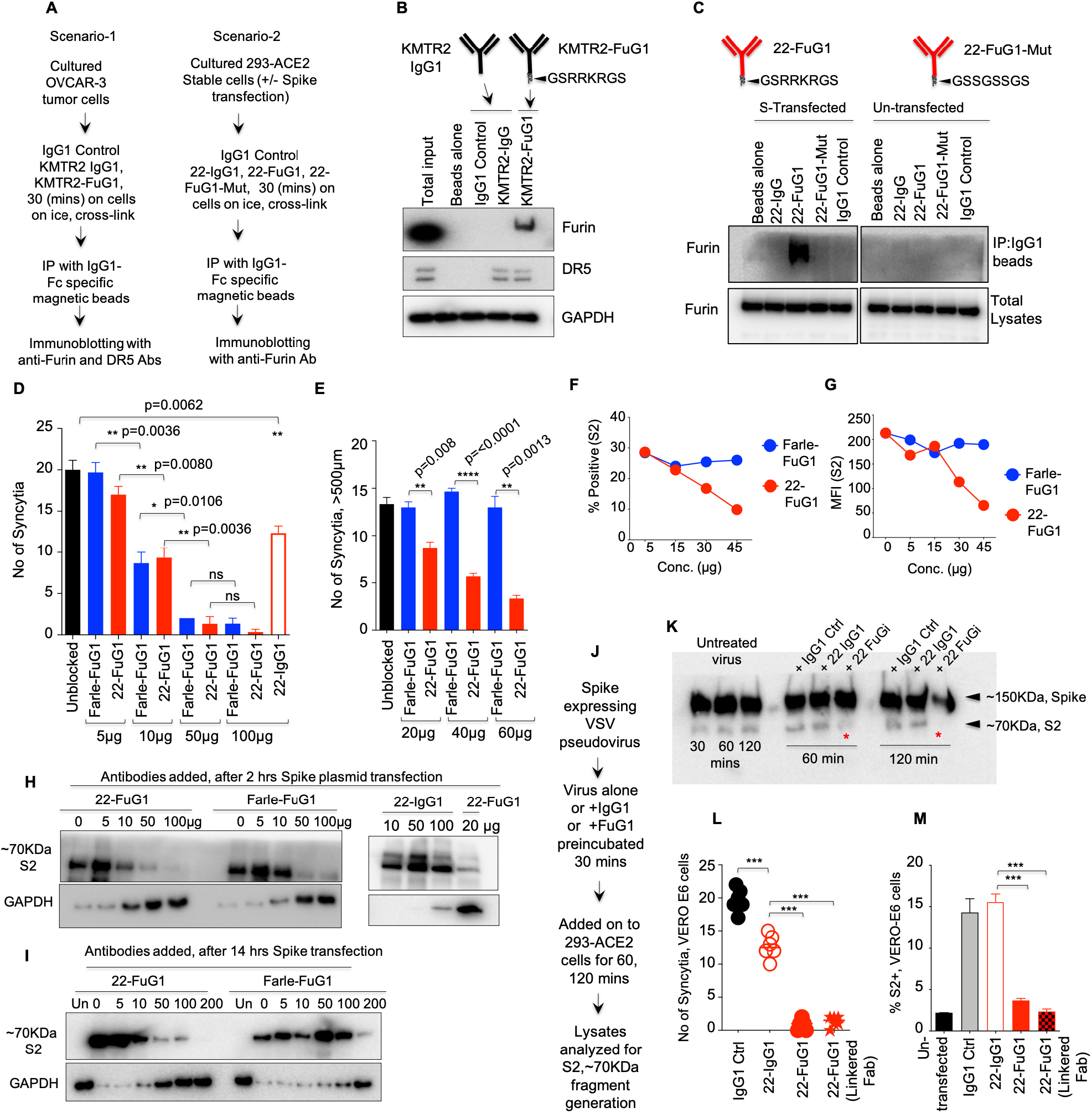
FuG1 antibody competitively interferes with the cellular furin. (A) Schematic of experiment shown in (B) and (C). (B) OVCAR-3 tumor cells were treated with indicated antibodies for 30 mins in cross-linking conditions. IP was carried out using IgG1-Fc specific magnetic beads followed by immunoblotting against DR5, furin. (C) Spike transfected (-/+) 293-ACE2 cells were treated with indicated antibodies. 22-FuG1-Mut lacks furin substrate residues in Fc-linker. IP was carried out similar to B. (D) Number of syncytia counted as shown in after indicated treatments (n=3). (E) Same as D, except indicated antibodies were added 14 hrs after transfection (n=3). (F-G) Same as E except, S2 generation was analyzed using flow cytometry. Data shows Mean S2+ (%) and MFI of cells after treatments (n=3). (H-I) Same as (D) and (E), excepts lysates were analyzed for generated S2 levels. (J-K) Pre-indication of spike pseudovirus inhibits spike-S2 conversion. (L-N) No. of syncytia and % surface S2 levels after indicated treatment using VERO-E6 cells.

To confirm 22-FuG1’s specificity against spike infected cells, we co-cultured comparable FOLR1^+^ 293-ACE2 and RFP-stable HCC-1806 cells (Fig. 3A). Spike transfected cocultures were treated with 50μg 22-IgG1, farletuzumab-FuG1 and 22-FuG1. As HCC-1806 are limitedly transfectable compared to 293 cells, we hypothesized equal FOLR1-dependent cell surface occupancy of farletuzumab-FuG1 against both cell-types in cocultures. On the contrary, 22-FuG1 would exclusively target spike^+^ cells (Fig. 3B). As expected, farletuzumab-FuG1 only reduced ~30% syncytia load compared to 22-FuG1 (~90%) in co-cultures (Fig. 3C-E) and only partially reduced S2 conversion in lysates (Fig. 3F). When tested in pseudovirus neutralization assays, 22-FuG1 strategy was significantly effective over RBD targeting IgG1, >6-fold higher infection neutralizing IC_50_ values (Fig. 3G-I).

**Figure 3.**
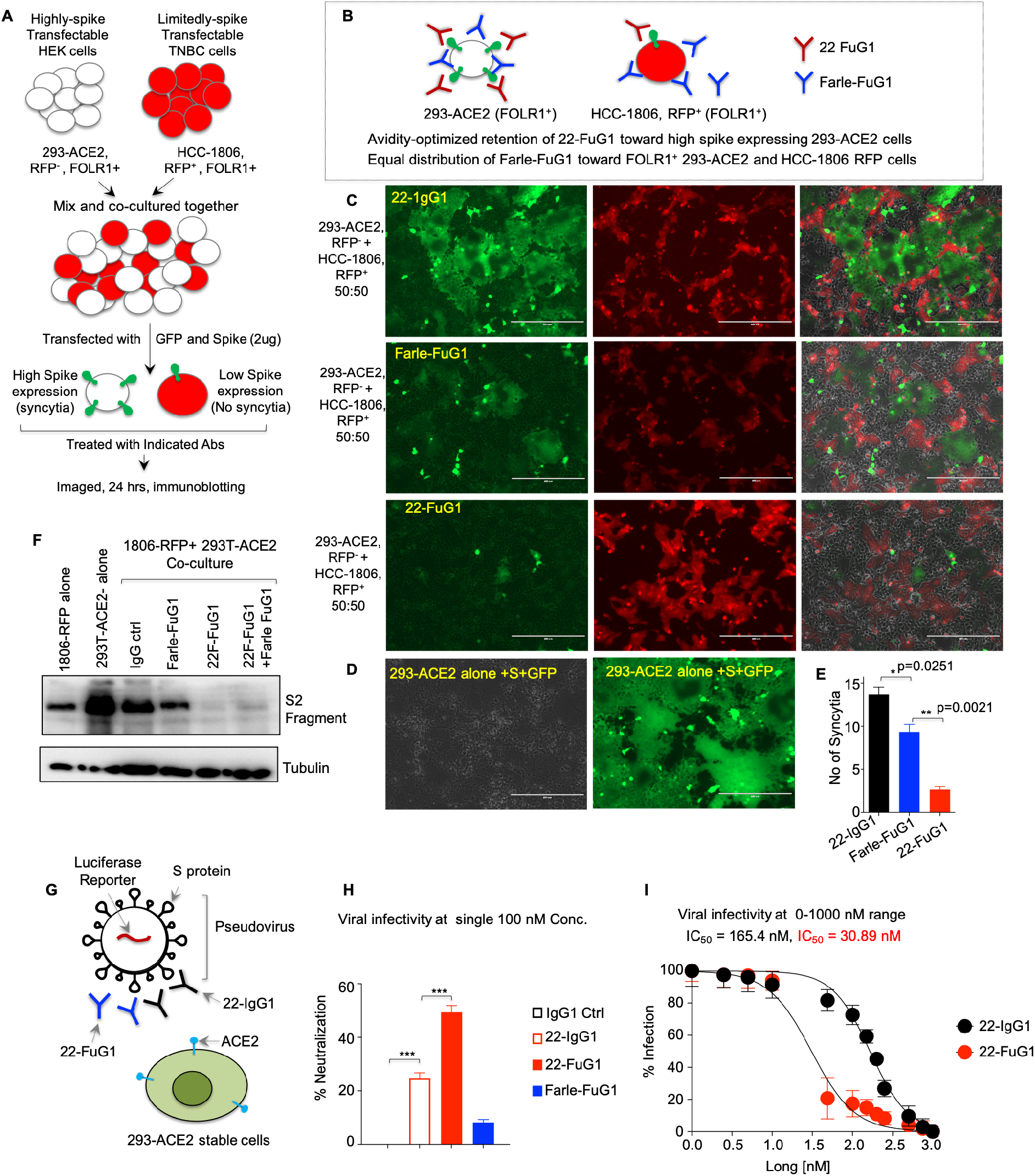
FuG1 antibody is selective in blocking SARS-CoV-2 entry. (A) Schematic of the co-culture experiment described in D and E. (B) Schematic showing preferred binding of 22-FuG1 (Red) to highly expressed spike on 293-ACE2 cells, while farletuzumab-FuG1 (Blue) distributes equally against both cell lines. (C-D) Co-cultured cells treated with 22-IgG1, Farle-FuG1 and 22-FuG1 antibodies. (E) Number of syncytia from C (n=3). (F) Total lysates from spike-transfected: HCC1806 cells, 293-ACE2 cells alone, and cocultured-cells treated with indicated antibodies were immunoblotted for S2. (G) Schematic of the experiment described in H, I. (H-I) Pseudoviral neutralization assays after treatment of Farle-FuG1 and 22-FuG1 antibodies at increasing concentration. IC_50_ values are shown (n=3).

## Conclusion

Our results describe an innovative plug-and-play strategy that targets the proteolytic mechanism of SARS-CoV-2 infection, the first step in viral pathogenesis. The described FuG1 strategy gives spike target mediated specificity over described furin and TMPRSS2 small molecule inhibitors(4). Furthermore, additional antibody dual-specificity based FuG1 co-targeting lung enriched receptor could be easily engineered on the described foundation for high specificity(6). As epitope of CR3022 does not fully overlap with ACE2-RBD binding(5, 8), FuG1 antibodies that strictly interfere with ACE2-RBD function are expected to be highly effective. In summary, we describe a rational peptide based therapeutic design of SARS-CoV-2 targetable strategy(9), that can be applied to newly identified SARS-CoV-2 variants(1) and the next generation of coronaviridae family posing threats to humans.

## Author contributions

Conceptualization, study design, Original Draft: J. T-S., Methodology: G.S., D.G., A.H, J.T-S., Supervision: J. T-S.

## Material and Methods

### Source of Antibody sequences

The sequence of Homo sapiens anti-SARS-CoV targeting immunoglobulin heavy chain and light chain are available at GenBank: DQ168569.1 (CR3022 VH) and GenBank: DQ168570.1 (CR3022 VL). The sequence of farletuzumab, KMTR2 etc. used are previously described(1) and are publicly available at IMTG.ORG

### Recombinant antibody Cloning

The sequence of various antibodies used in this study is provided in Table S1. All antibodies were clones and expressed as described earlier(1). Briefly, the DNA sequences were retrieved from the open sources (IMGT.ORG or publicly available patents or NCBI etc.) and synthesized as gene string with overlapping region to pCDNA3.1 vector using Invitrogen GeneArt gene synthesis services. Using overlapping pCDNA3.1 restriction site EcoR1 and HindIII primers, PCR amplification was carried out of gene VH and VL gene strings separately. After PCR amplification, DNA was gel purified and inserted into pCDNA 3.1 vectors (CMV promoter) by making use of In-Fusion HD Cloning Kits (Takara Bio). EcoR1 and HindIII digested vector was incubated with overlapping PCR fragments (of various different recombinant DNAs, see list of clones in Key Resource Table) with infusion enzyme (1:2, vector: insert ratio) at 55°C for 30 min, followed by additional 30 min incubation on ice after adding *E. coli* Stellar™ cells (Clontech, see reagent Table). Transformation and bacterial screening were carried out using standard cloning methods. Positive clones were sequenced confirmed in a 3-tier method. Confirmed bacterial colonies were Sanger sequencing upon PCR followed by re-sequencing of mini-prep DNA extracted from the positive colonies. Finally, maxiprep were re-sequenced prior to each transfection. Recombinant antibodies were also re-confirmed by ELISA and flow cytometry surface binding studies as described earlier(1). FuG1 antibodies were also engineered in similar way. A standard furin substrate site is provided for Fc-extended linker sequence in manuscript and the optimal sequence is part of our intellectual proprietary info here at UVA. Linkered monospecific and bispecific with knob and hole mutations were generated using flexible linkers as described previously (1).

### Recombinant RBD IgG4-Fc cloning

To clone CR3022 binding RBD domain, amino acid 333-530 were order for gene string synthesis in continuation of G4S flexible linker and IgG4 CH2 and CH3 domain with EcoR1 and HindIII overlapping region. Following PCR amplification with overlapping primers, DNA was gel purified and inserted into pCDNA 3.1 vectors (CMV promoter) by making use of In-Fusion HD Cloning Kits (Takara Bio) as described in previous section. Recombinantr IgG4-FC FOLR1 and DR5 were also cloned in similar manner and have been described previously (1).

### Recombinant antibody and IgG4-Fc antigen expression

Free style CHO-S cells (Invitrogen, Reagents Table) were cultured and maintained according to supplier’s recommendations (Life technologies) biologics using free style CHO expression system (life technologies) and as previously described(1, 2). A ratio of 1:2 (light chain, VL: heavy chain, VH) DNA was transfected using 1μg/ml polyethyleniamine (PEI, see Key Resource Table). After transfection cells were kept at 37°C for 24 hrs. After 24 hrs, transfected cells were shifted to 32°C to slow down the growth for 9 additional days. Cells were routinely feed (every 2^nd^ day) with 1:1 ratio of Tryptone feed and CHO Feed B (see Reagents Table). After 10 days, supernatant from cultures was harvested and antibodies were purified using protein-A affinity columns. Various recombinant antibodies used in this study and recombinant target antigens were engineered, expressed and purified in Singh Laboratory of Novel Biologics as described earlier(1). Recmbinants antigens were similar expressed and transfected with PEI at 1 μg/ml concentration. All prufied proteins were also confirmed for size using reducing SDS-PAGE, and standard ELISA as described earlier(1).

### Antibody purification

Various transfected IgG1 antibodies, FuG1 antibodies and recombinant IgG4-Fc antigens (as indicated in text and figure legends) were affinity purified using HiTrap MabSelect SuRe (GE, 11003493) protein-A columns. Transfected cultures were harvested after 10 days and filtered through 0.2-micron PES membrane filters (Milipore Express Plus). Cleaning-in-place (CIP) was performed for each column using 0.2M NaOH wash (20 min). Following cleaning, columns were washed 3-times with Binding buffer (20 mM sodium phosphate, 0.15 M NaCl, pH 7.2). Filtered supernatant containing recombinant antibodies or antigens were passed through the columns at 4°C. Prior to elution in 0.1M sodium citrate, pH 3.0-3.6, the columns were washed 3 times with binding buffer (pH 7.0). The pH of eluted antibodies was immediately neutralized using sodium acetate (3M, pH 9.0). After protein measurements at 280 nm, antibodies were dialyzed in PBS using Slide-A-Lyzer 3.5K (Thermo Scientific, 66330). Antibodies were run on gel filtration columns (next section) to analyze the percent monomers. Whenever necessary a second step size exclusion chromatography (SEC) was performed. Recombinants IgG4-Fc tagged extracellular domain antigens such as rFOLR1, rDR5 and RBD etc. were also similarly harvested and purified using protein-A columns.

### Size Exclusion chromatography

The percent monomer of purified antibodies was determined by size exclusion chromatography. 0.1mg of purified antibody was injected into the AKTA protein purification system (GE Healthcare Life Sciences) and protein fractions were separated using a Superdex 200 10/300 column (GE Healthcare Life Sciences) with 50mM Tris (pH 7.5) and 150mM NaCl. The elution profile was exported as Excel file and chromatogram was developed. The protein sizes were determined by comparing the elution profile with the gel filtration standard (BioRad 151-1901) (Hong et al., 2012). Any protein peak observed in void fraction was considered as antibody aggregate. The area under the curve was calculated for each peak and a relative percent monomer fraction was determined as described earlier (1)

### Binding studies by ELISA

Binding specificity and affinity of various described IgG1’s and FuG1 antibodies (including linkered FuG1, FuG1-Lin) s were determined by ELISA using the recombinant extracellular domain of corresponding receptor/target antigen. For coating 96-well ELISA plates (Olympus), the protein solutions (2μg/ml) were prepared in coating buffer (100mM Sodium Bicarbonate pH 9.2) and 100μl was distributed in each well. The plates were then incubated overnight at 4°C. Next day, the unbound areas were blocked by cell culture media containing 10% FBS, 1% BSA and 0.5% sodium azide for 2 hr at room temperature. The serial dilutions of antibodies (2-fold dilution from 50 nM to 0.048 nM) were prepared in blocking solution and incubated in target protein coated plates for 1 hr at 37°C. After washing with PBS solution containing 0.1% Tween20, the plates were incubated for 1 hr with horseradish peroxidase-(HRP) conjugated anti-human IgG1 (Thermo Scientific, A10648). Detection was performed using a two-component peroxidase substrate kit (BD biosciences) and the reaction was stopped with the addition of 2N Sulfuric acid. Absorbance at 450 nm was immediately recorded using a Synergy Spectrophotometer (BioTech), and background absorbance from negative control samples was subtracted. The antibody affinities (Kd) were calculated by non-linear regression analysis using GraphPad Prism software.

### Binding studies by BioLayer Interferometry (BLI)

Binding measurements were performed using Bio-Layer Interferometry on FortéBio Red Octet 96 instrument (Pall) as described earlier(1). Biotin-Streptavidin based sensors were employed for the studies. Recombinant Fc linked DR5 variants were biotinylated using EZ-Link Sulfo-NHS-SS-Biotin (Thermo Scientific 21331) following the manufacturer’s instructions. Unreacted Sulfo-NHS-SS-Biotin reaction was quenched by 50mM Tris-Cl pH 7.4 and removed via dialyzing against PBS. For binding analysis biotinylated antigen (1 μg/mL) were immobilized on streptavidin (SA) biosensors (Pall) for 300 sec to ensure saturation. Associate and dissociation reactions were set in 96-well microplates filled with 200 μL of unbiotinylated DR5 agonist for 900 Sec. All interactions were conducted at 37 °C in PBS buffer containing 2mg/ml BSA. These binding observations were also confirmed by biotinylating the agonist antibodies and probed against unbiotinylated DR5-Fc variants. Kinetic parameters (K_ON_ and K_OFF_) and affinities (KD) were analyzed using Octet data analysis software, version 9.0 (Pall).

### Western Blotting

Cells were cultured overnight in tissue culture-treated 6-well plates prior to treatment. After antibody treatment for 48 hr (or indicated time), cells were rinsed with PBS and then lysed with RIPA buffer supplemented with protease inhibitor cocktail (Thermo Scientific). Spinning at 14000 rpm for 30 min cleared Lysates and protein was quantified by Pierce BCA protein assay kit. Western blotting was performed using the Bio-Rad SDS-PAGE Gel system. Briefly, 30 μg of protein was resolved on 10% Bis-Tris gels and then transferred onto PVDF membrane. Membranes were blocked for one hour at room temperature in TBS + 0.1% Tween (TBST) with 5% non-fat dry milk. Membranes were probed overnight at 4°C with primary antibodies. Membranes were washed three times in TBST and then incubated with anti-rabbit or anti-mouse secondary antibodies (1/10,000 dilution, coupled to peroxidase) for 1 hr at room temperature. Membranes were then washed three times with TBST and Immunocomplexes were detected with SuperSignal West Pico Chemiluminescent Substrate (Thermo Fisher Scientific). Images were taken using a Bio-Rad Gel Doc Imager system. Primary antibodies are listed in the Key Resource Table.

### Syncytia formation and inhibition

293T cells expressing stable ACE2 (293-ACE2) have been described earlier(3) and were kindly provided Dr. Jesse.D. Bloom. 293-ACE2 were transfected with 1μg (unless mentioned) of Spike GFP plasmid using Mirus reagent as per manufacturer’s instructions. Transfection media was replaced with fresh media after 4hr of transfection. Cells were monitored for the formation of syncytia after 20h of incubation. In case of syncytia inhibition experiment, various control and FuG1 antibodies were added with indicated concentrations (see figures and figure legends for specific concentration) in different set of experiments. For some experiments, control and FuG1 antibodies were added 2 hrs after transfection and for other 8 hrs after transfection to allow effective spike cellular expression (as indicated in figures and figure legends).

### Flow cytometry

The cell surface expression of ACE2, Spike, DR5, FOLR1 etc. was analyzed by flow cytometry in various experiments (see figure and figure legends) after treatment with various control and FuG1 antibodies. After various treatments, cells were trypsinized and suspended in FACS buffer (PBS containing 2% FBS). The single cell suspension was then incubated with primary antibodies for 1 hr at 4°C with gentle mixing. Following wash with FACS buffer, the cells were then incubated with fluorescently labeled anti-Rabbit antibody for 1 hr. Cells were washed and flow cytometry was performed using FACSCalibur. The data was analyzed by FCS Express (De Novo Software) and FlowJo.

### Production of SARS-CoV-2 S pseudovirions

Pseudovirions were generated as described previously(4). Briefly, BHK21 cells were seeded at a density of ~2×10^6^ cells per well into a 24-well cell culture plate in 500 μl of DMEM media. Cells were incubated at 37°C with 5% CO2 overnight. Media was aspirate and cells were co-transfected with pCAAG Spike DNA (0.5 μg) and 7 μl of VSV-G pseudovirus in 500 μl of 1X opti-MEM reduced serum free media using TransIT-Pro^®^ as a transfection reagent (Mirus). After 4 hr of transfection, Opti-Mem media was changed to regular DMEM media and incubated at 37°C with 5% CO2. Two days post transfection; supernatants were collected and clarified by centrifugation at 1000 rpm for 5 min, passed through 0.45 μm PES filter, and concentrated using 10% (w/v) PEG 6000 and 5M NaCl. PEGylation solution was mix and incubated at 4°C overnight on gentle rocker. The precipitated pseudovirus particles were centrifuged at 4000 rpm for 30 min and resuspended in 1 ml of culture media and aliquoted prior to storage in −80°C to avoid multiple freeze-thaws. To transduce cells with pseudovirions, ~2 x10^5^ 293T cells were seeded in a 96 well plate 20-24hrs prior to infection. A single aliquot of pseudo-virus was used for infection and titrations were performed in technical triplicates. After overnight incubation, cells were fed with fresh media. About 48 h post transduction, 5 μl D-luciferin (300 μg/ml) reagent was added per well and incubated at room temperature for 15 minutes. The transduction efficiency was measured by quantification of the luciferase activity using a Synergy HT 96 well microplate reader (BioTek Instruments Inc, USA). All experiments were done in triplicates and repeated at least twice or more.

### Pseudovirus neutralization assay

For viral neutralization, we confirmed the experiments with commercially purchased pseudoviral particles from two different suppliers, BPS Biosciences (SARS-CoV-2 Spike Pseudotyped Lentivirus with Luciferase Reporter) and Virongy (SARS-CoV-2 (Luc) Pseudovirus), See Key Resource table for more details. Furthermore, we also generated pseudoviral particles in our lab as described previously (4, 5). Briefly, 293T-ACE2 cells were grown in 96 well plate. SARS-CoV-2 Spike Pseudotyped Lentivirus containing Luciferase Reporter were purchased and tested with laboratory generated viral particles side-by-side. The viral neutralization experiment was set up with 2μl of virus (3×10^5^ TU/ml) for BPS Biosciences pseudovirus (Cat # 79942) and 10μl of virus for Virongy pseudovirus as per manufacture recommendations. 100nM control and FuG1 antibodies were used in viral neutralization assay (Luciferase readout) described in Figure 3I. 0.1-1000nM concertrations of antibodies were were used in viral neutralization assay (Luciferase readout) described in Figure 3J. Following incubation at room temperature for 30min, the reaction mixture was added to the cells. Plates were incubated for 48-60h at 37C. All experiments were done in triplicates, and repeated at least thrice or more. The viral infectivity was measured by quantifying luciferase activity using 150ug/ml of XenoLight D-Luciferin bioluminescent substrate (PerkinElmer # 122799). Data was plotted using GraphPad Prism software.

### Determination of Inhibitory concentration (IC_50_ Value)

For inhibitory concentration determination, different concentrations (0-1000 nM) of antibodies (22-IgG, 22-FuGi and IgG1 control) were pre-incubated with 7 μl of SARS-CoV-2 (Luc) lenti-pseudovirus in a total volume of 10 μl for 30 minutes. After incubation, virus/antibody mixed was added onto ACE2-HEK293 cells. For control wells, the same numbers of ACE2-HEK293 cells were seeded, but no antibody was added. Approximately, 48-60 hours after transduction, percent infection was calculated by measuring luciferase activity. The infection caused by the virus only (without antibody treatment) was considered to be 100% activity and the decrease in percent infection was compared to that and IC_50_ value was calculated using GraphPad Prism software. All experiments were done in triplicates, and repeated at five times.

### Quantitation and statistical analysis

Data unless indicated otherwise are presented as mean ±SD. In general, when technical replicates were shown for *in vitro* experiments, student t-test was used for statistical analysis and the same experiment was at least repeated once with similar trend observed. When data from multiple experiments was merged into one figure, statistical significance was determined by an Wilcoxon Mann-Whitney test using Graph Pad Prism 5.0 software. For all the statistical experiments p values, p<0.05 (*), p<0.01 (**) and p<0.001 (***) were considered statistically different or specific p values indicated otherwise or “ns” indicates non-significant.

